# Quantifying the thermostability landscape separating plant sesquiterpene synthase orthologs using Maximum Entropy

**DOI:** 10.1101/172437

**Authors:** Charisse M. Nartey, Johnatan Aljadeff, Irma Fernandez, Caroline Laurendon, Hyun Jo Koo, Paul E. O’Maille, Tatyana O. Sharpee, Joseph P. Noel

**Author notes:** Co-first author. For correspondence (TOS); (JPN).

## Abstract

As global temperatures rise due to climate change, crops are becoming increasingly vulnerable to failure. We tested the accessibility of thermodynamic phenotypic diversity within limited positional changes using 2^9^=512 mutants of a terpene synthase (TPS), Tobacco 5-epi-Aristolchene synthase. First, we measured the thermal unfolding curves of each mutant and found that mutations shifted the T_m_s both higher and lower, including a cohort of mutants that failed to fold. The low correlation coefficient between the T_m_s of these mutants and the proportions of each terpene product output by the enzymes revealed that thermostability and product output are independent traits. Maximum Noise Entropy analyses were used to analyze the impact of the 9 mutational positions on thermostability, revealing that three of these positions were mainly responsible for the trait of increased T_m_. These positions form a functional network as measured by the nonlinearity of their combined effects on thermostability. Unexpectedly, the strongly destabilizing positions combine nonlinearly to ameliorate each other’s deleterious effects resulting in a synergistic dampening. Taken together, our study shows the high potential for specialized metabolic enzyme engineering but also reveals a complex interconnected system of amino acids that will continue to evade perfect predictability.

---

Sustainable and climate change-resistant crops are not only critical for future food security but also global geopolitical stability^1^. Harnessing the combined power of protein engineering and breeding techniques will continue to aid farmers in avoiding major crop failures due to regional warming, disease and environmental stresses such as increases in consistent water deficits^2,3^. In addition, these strategies may improve crop yields to meet the needs of a global population that is predicted to increase by 33% over the next 34 years^4^.

While the techniques for genetically modifying crop species continue to expand^5^, further research is needed to uncover first principles that might accelerate protein engineering approaches. To address this growing need, we focus on specialized metabolic enzymes, as their prominent role in plant ecology and large diverse gene families make them key players in the push to enhance crop yield and robustness as average temperatures rise.

Tobacco (*Nicotiana tabacum*) EAS (TEAS) and Henbane (*Nicotiana tabacum*) premnaspirodiene synthase (HPS) are encoded by homologous genes (∼80% DNA exon sequence identity), yet cyclize the ionized form of the acyclic primary metabolite farnesyl diphosphate (FPP) into bicyclic and chiral 5-epi-aristolochene (5-EA) and premnaspirodiene (PSD), respectively, both of which serve as precursors to species-specific antimicrobial phytoalexins (Fig. 1a). In general, TPSs often exhibit broad product profiles that vary with specific mutational differences, temperature and pH^6^. TEAS generates at least 24 minor products in addition to 5-EA^7^. Thus, achieving a predictive understanding of mutational pathways leading to new fitness attributes not only has practical applications to ecology and plant sustainability in light of climate change^8^ but also to the flavor and fragrance industry and to drug discovery and improvement^9,10^.

**Fig. 1.**
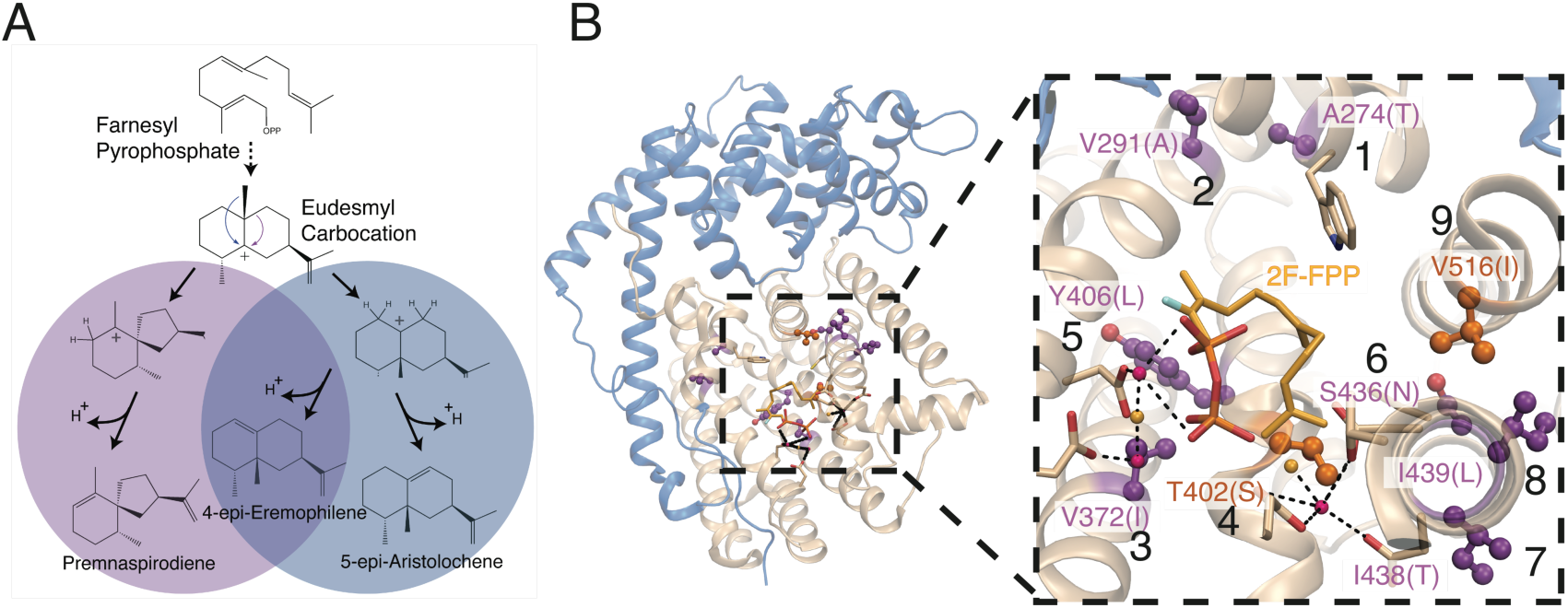
Structural overview of TEAS catalysis and M9 library positions. **(A)** Catalytic pathways generating the major products of TEAS library mutants. (**B**) Crystal structure of WT TEAS bound to 2-fluoro-FPP (PDB ID: 3M02). M9 positions are highlighted in orange or purple based on their direct or indirect contact with active site surface, respectively. Positions are labeled from 1-9 according to their sequence order and are hereafter referred to in the text by this convention. Structural images made in Pymol.

Several chemical steps in the catalytic mechanism of TEAS are shared with HPS. We previously demonstrated that by exchanging 9 amino acids in and surrounding the active sites of wild type (WT) TEAS and WT HPS generating M9 TEAS and M9 HPS, respectively, we reassigned and interconverted their product profiles without significantly compromising catalytic efficiencies^11^. Further, we generated, expressed, and biochemically analyzed a TEAS mutant library spanning all possible combinations of these 9 naturally mutable positions (numbered 1-9, giving 2^9^ = 512 mutants. See Fig. 1b) demonstrating that protein epistasis, or nonlinearity, modulates catalytic specificities of TEAS mutants^12^.

To quantify the extent and magnitude of protein epistasis vis-à-vis thermostability across all 9 positions, we expressed, purified and measured the thermostability of all 512 TEAS M9 library mutants. While an *in vitro* examination of thermostability does not incorporate the role of chaperones and other factors important for folding, it does provide quantitatively-rich measurements of this adaptable biophysical property. We then applied computational tools, including Maximum Noise Entropy^13-19^, to extract statistically significant patterns of amino acid residue contributions to variations in the thermal unfolding profiles (e.g. curve shapes), including changes to the temperature midpoint of unfolding (T_m_). These analyses made it possible to rank the contribution of each of the 9 mutated positions to the characteristic shapes of the unfolding curves and the thermostabilities quantified by T_m_. Across the M9 library, phenotypes cluster around WT-like thermostability, with half of the positions observed to stabilize the protein while the other half destabilize the protein relative to WT TEAS. Importantly, we discovered, that on average, pairs of these residues dampen each other’s individual effects on thermostability. This set of observations suggests that counterbalancing effects of outer tier active site mutations enable TEAS mutants to straddle the line between remaining mutationally robust (stable) yet amenable to catalytic adaptation and selection from a population of divergent sequences and traits.

## Results

### Thermostability phenotypes

To assess the temperature-dependent unfolding properties of the 512-member M9 TEAS mutant library, we overexpressed proteins in *E. coli*, purified the soluble fraction via Ni^2+^-NTA affinity chromatography, and quantified the purity and concentration of samples by SDS-PAGE and band densitometry of Coomassie stained gels. We then measured the thermal unfolding of purified protein samples via thermofluor assays, which monitor the extent of protein unfolding using fluorescence changes of an environmentally sensitive non-polar dye as a function of temperature ^20^ (Fig. S1a). The changes in slopes of the change in fluorescence versus temperature curves correlate with the cooperativity of unfolding of the protein ensembles, and therefore, often reveal the extent of sample conformational homogeneity during unfolding ^21^. Since the concentrations of soluble protein varied between ∼0-5 uM, we normalized the fluorescence data traces (see Methods). We observed a range of thermal stability phenotypes across the M9 library with T_m_ shifts ranging from −4.1 to +11.9 °C relative to WT TEAS.

In order to compare all curves without bias to particular temperatures, we calculated the geometric distances between each pair of profiles, thereby generating a 512X512 distance matrix (Fig. S1b). We used this matrix as the input to compute an unrooted neighbor-joining “thermostability tree” (Fig. 2). This tree displays the wide diversity of phenotypes accessed within the sequence space spanned by the TEAS M9 library, and illustrates the degree to which specific sequences cluster to form clades and families in much the same way as previously observed for the product profiles of the M9 library ^12^. We annotated each clade with the superimposed thermofluor data from each mutant represented therein (grey curves) and included the average of these curves to highlight the characteristic shape of each clade (colored curves). Each of these plots is also labeled with the average T_m_s of the included mutants as well as the average concentration of purified protein used for each set of measurements.

**Figure 2.**
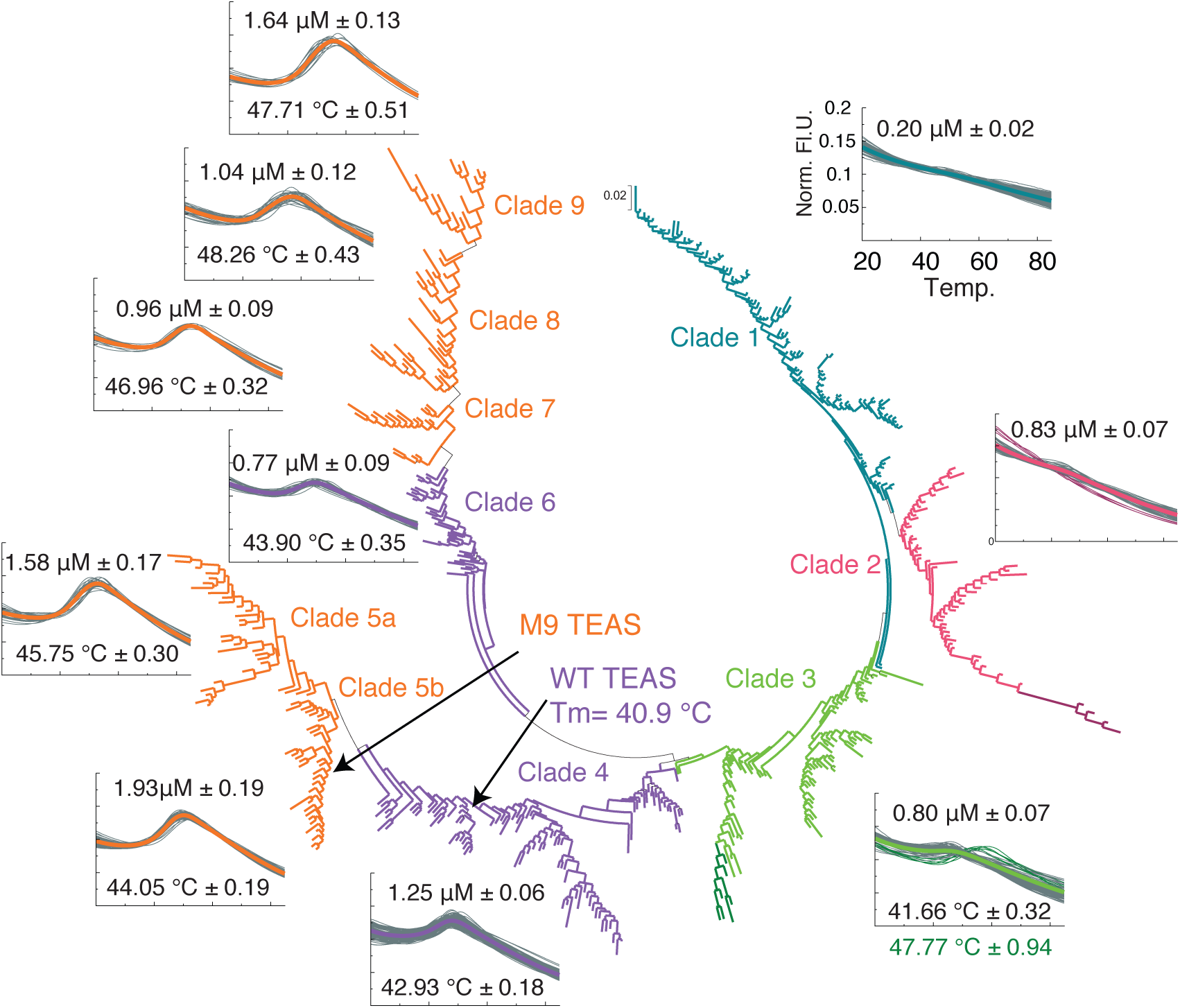
Thermal unfolding phenotypes reveal spread of mutational effects on thermostability in the TEAS M9 library. Neighbor-joining tree of thermal unfolding phenotypes in the TEAS M9 protein library. Raw fluorescence data were averaged (4 replicates), condensed into 100 bins, normalized such that the norm of each curve equals 1, and a 512x512 distance matrix calculated. The neighbor-joining tree was generated from these distances using Mega 5.2 ^78^. Clades were assigned based on tree topology and grouped into phenotypic families by average distance between clades. X-and Y-axes are temperature in °C and Norm Fl.U, respectively. Each clade is annotated with the superimposed thermofluor data from each mutant represented therein (grey curves) and the average of these curves highlights the characteristic profile of each clade (colored curves). Each plot is also labeled with the average T_m_s of the included mutants as well as the average concentration of purified protein used for each set of measurements. The range of T_m_ across the library is 36.8 −52.8 °C. Many unfolding curves have unfolding transitions that are too shallow to obtain a reliable T_m_ either because the protein concentration is too low or because the sample is already significantly unfolded. In Clade 2, a few mutants diverge significantly from the average (highlighted in burgundy), and in Clade 3, there are 7 standout mutants with higher T_m_ than WT (highlighted in dark green). Averaged T_m_ and average recovered protein concentrations for mutants are displayed. Average T_m_ for Clade 3 is calculated for 54/89 mutants, which excludes the dark green, standout mutants and the mutants for which a T_m_ could not be measured accurately. All errors are standard error of the mean.

Since the WT control resides in Clade 4, we designate this cluster and its closest neighbor (in terms of inter-clade distance), Clade 6, the WT-like family (purple clades). Many mutants in clades 5 and 7-9 are characterized by elevated T_m_s relative to WT (orange clades, Fig. 2); several mutants unfold at greater than 50 °C, while the T_m_ of WT is 40.9°. Clades 1 and 2 are populated by mutants having weak to nonexistent unfolding transitions consistent with two likely properties. One, the protein samples may exhibit considerable dynamics associated with unfolding transitions at room temperature. Two, the concentration of soluble samples recovered from heterologous expression and purification are low (see concentrations in Fig. 2). Normalization of the thermofluor dataset renders these two phenotypes essentially indistinguishable. Many mutants in clade 3 also display modest unfolding transitions with modest fluorescence increases, for a combination of the same reasons; nevertheless, 54 of the 89 mutants in clade 3 had measureable T_m_s, many of which are up to 3 °C lower than WT TEAS.

Strong interdependence within biophysical phenotypes is a major impediment to facile protein engineering. If a specific mutation results in both Tm changes and product profile changes, then we cannot enhance thermal robustness without potentially damaging the catalytic properties and vice versa. To address whether this type of obstacle exists in our system, we calculated the correlation coefficients between the 12 terpene products and the Tm values of each mutant for which a Tm could be detected by the Light Cycler software. As discussed above, mutants from Clades 1 and 2 and some from Clade 3 have weak fluorescence profiles and were excluded from the analysis, leaving 313 mutants total for the calculation. As shown in Figure S1, there is no correlation between thermostability and any of the products. Taken together these results illustrate that thermodynamically robust phenotypes are accessible through changes at a relatively few numbers of sites in the TPS fold and that two important biophysical properties of these enzymes, product profile and thermostability, can be optimized independently of one another.

### Maximum Noise Entropy (MNE) quantifies phenotypic contributions of each mutation

To directly characterize how the presence or absence of a given mutation affects the thermal unfolding profiles of sequence populations, we used the Maximum Noise Entropy (MNE) method, previously applied to identify relevant patterns in neuroscience datasets ^18,19,22^. This method determines patterns that best separate the set of fluorescence curves that are associated with the presence of a particular mutation (representing 50% of library mutants) from the set of curves associated with the absence of this mutation across the library of mutants. This method, akin to a logistic regression, works despite the non-Gaussian and correlated structure of thermal unfolding profiles ^18,19^. The results of the analyses consist of linear combinations of fluorescence changes in the range of 20-85°C that are most strongly associated with mutation at a particular amino acid position (Fig. 3a). Notably, these quantities cannot be computed by direct averaging of raw unfolding curves.

**Figure 3.**
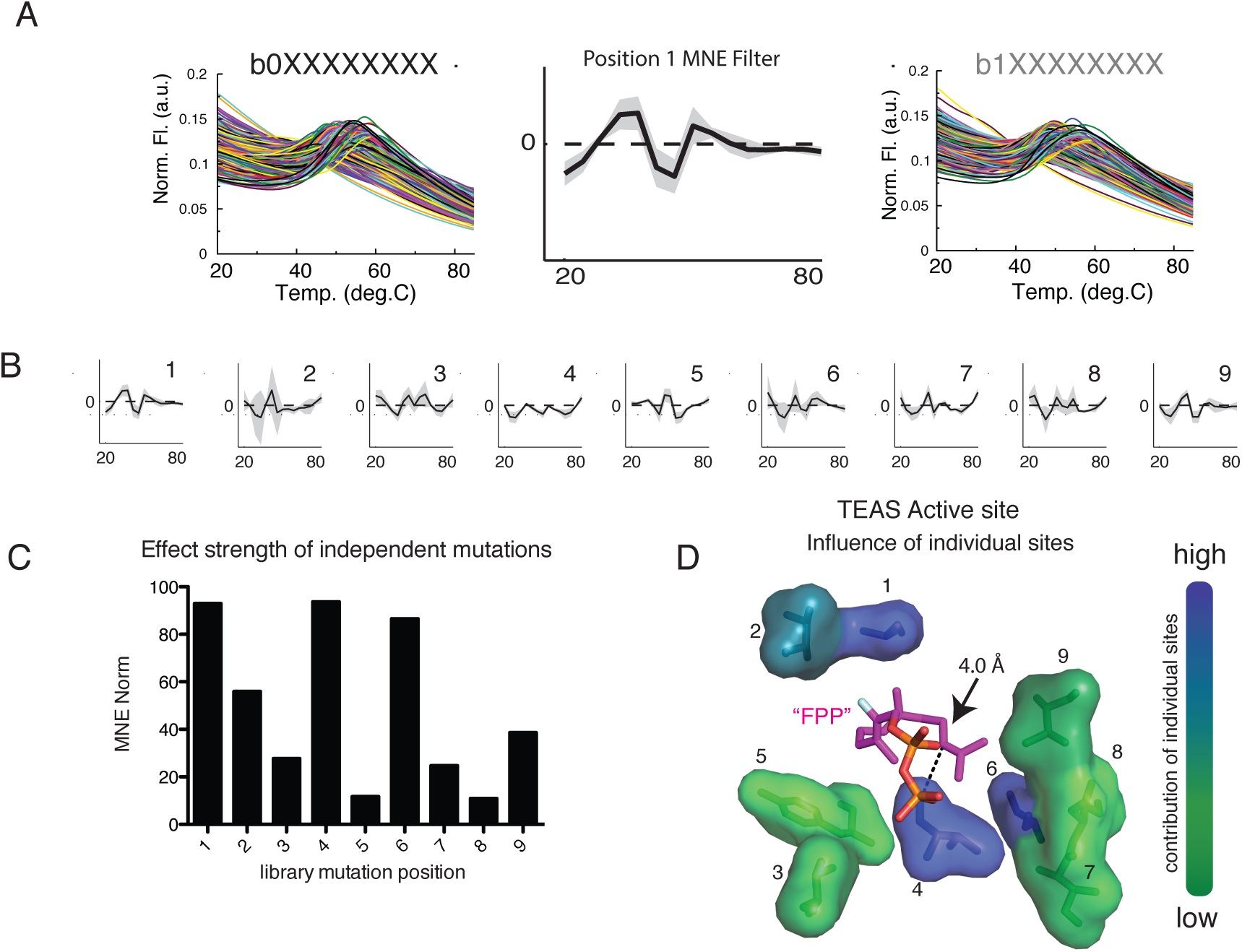
Maximum Noise Entropy Analysis of Thermofluor data. (A) Schematic representation of a first order MNE profile acting as a mathematical filter for separating two populations of unfolding traces with maximal probability. (B) Normalized MNE profiles capture the first order effect of each of the nine mutation sites. Gray shaded areas represent confidence bounds found by separating the data into training and validation sets. All MNE profiles were found to be significant based on the KL distance (Cover & Thomas 1991) between the distribution of projection onto the relevant dimension and the same distribution computed for randomly assigned mutations. This tests aims to compare significance relative to the cases where there is no association between mutations and changes in the thermal unfolding profile. (C) The strength of each library mutation is proportional to the KL distance between distributions of projections of the MNE profile onto the two populations of thermofluor curves. (D) The 9 library positions are displayed in the 3D active site structure of WT TEAS (PDB ID: 3M01) with a substrate analog 2-fluoro-farnesyldiphosphate (2F-FPP) bound (grey sticks). The library amino acids are rendered as sticks covered by a 20% transparent van der Waals surface representation. A color gradient from green to blue indicates the norm with values ranging from ∼11-94. Ray tracing fog feature was removed to ensure accuracy of color gradient.

The relative strength of the mutation’s effects, which we will hereafter refer to as the “intensity” of the MNE profile, is represented by the magnitude (technically L_2_ norm) of the computed MNE profile for this particular mutant’s population of sequences (see Extended Experimental procedures for further details). MNE profiles for the 9 library positions calculated individually are shown in Figure 3b. Overall, each amino acid position exerts statistically significant effects (see Ext. Experimental Procedures) on the thermostabilities of the position’s mutant populations. When ordered by the intensity of the MNE profile, the positions formed the sequence 4, 1, 6, 2, 9, 3, 7, 5, 8 from largest to smallest MNE intensities (Fig. 3c). The effect each mutation has on changes to fluorescence varies in complex ways between positive and negative values across the temperature dimension, confirming the existence of thermostability features influencing fluorescence changes at temperatures far from the WT TEAS T_m_. Mapping the intensity values of these MNE models onto the 3D structure of the TEAS active site reveals that the most influential positions, 4, 1 and 6, are spread across the active site, and are distal from each other rather than forming a spatially connected core (Fig. 3d).

### Biophysical interpretation of MNE profiles

To quantify the expected effect of each mutation in terms of T_m_ changes, we compared MNE profiles for each mutation with those expected for a pure shift in T_m_ compared to WT TEAS with no other changes to the WT TEAS unfolding profile. We refer to these analyses as Basis Function Analyses. Mathematically, given the WT thermal unfolding profile *F*(*T*), if a site-specific mutation only modifies T_m_ but does not otherwise affect the shape of the unfolding curve, then the expected change in the unfolding profile will occur along the derivative of the WT unfolding profile, *F*′(*T*) (Fig. 4a). One can then estimate the magnitude of the T_m_ shift by comparing the MNE profile to the appropriately normalized derivative of the WT unfolding profile with respect to temperature changes, *ΔT*, to obtain dT_m_ in units of °C (see Experimental Procedures and Extended Exp. Proc.). These Basis Function analyses extract portions of the MNE profiles that describe T_m_ shifts relative to WT TEAS. Figure 4b illustrates these results for the single mutant populations; listed in order of decreasing effects, positions 5, 2 and 8 are stabilizing with regard to increased T_m_s, while positions 6, 1 and 3 are destabilizing. Here, we find position 4 has only modest effects on T_m_ shifts. This position, which exerts the greatest influence on variation within the thermal unfolding dataset, primarily results in changes in the shapes of thermal unfolding curves that do not correlate with changes to T_m_s.

**Figure 4.**
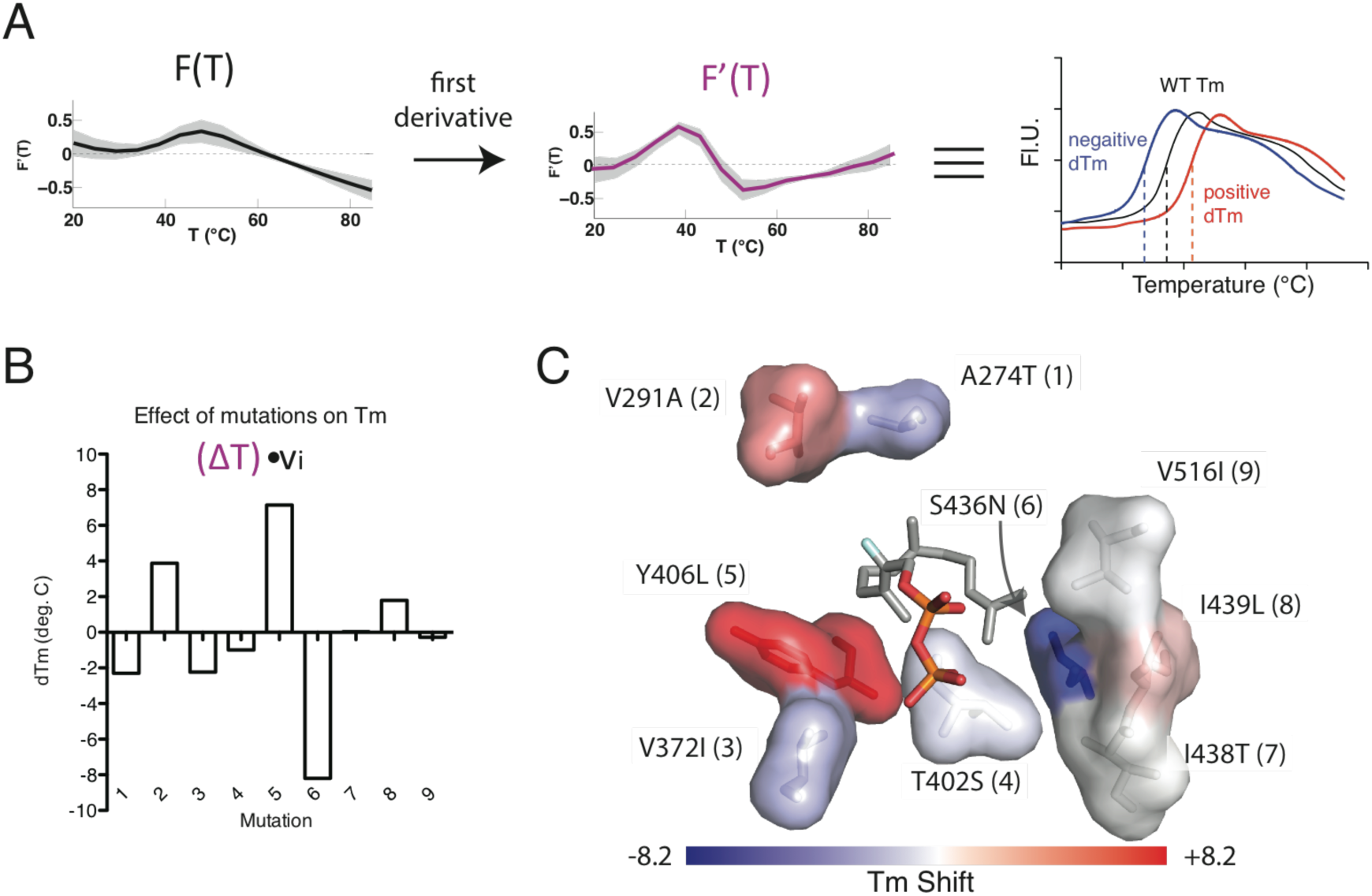
Basis Function Analysis to extract T_m_ shifts from MNE profiles. A. The basis function describing Tm shifts is the first derivative of the WT unfolding curve *F*(*T*). This function is then multiplied by the covariance matrix of the binned, normalized fluorescence dataset to obtain ?T. (B) The projection of this vector onto the MNE profiles gives the T_m_ shift in °C. (C) The dTm values are plotted onto the 3D structure (using PDB ID 3M01), which shows that pairs of interacting residues have opposing effects on T_m_. Blue to white to red gradient represents T_m_ shifts from −8.2 to 0 to 8.2 °C.

Previous studies have shown that amino acid positions that affect similar functions of the enzyme cluster in the 3D protein structure to form groups called “sectors” ^23,24^. Such an arrangement would enhance and expedite engineering efforts, because educated guesses can be made regarding the role of uncharacterized positions that are nearby well-characterized positions. To test whether this is the case in our system, we mapped the MNE-derived T_m_ values for all nine positions onto the 3D structure of the TEAS active site (Fig. 4C). Instead of spatial clustering of stabilizing and destabilizing mutations, respectively, surprisingly we found their distribution in 3D space to be statistically indistinguishable from random chance. This result shows that the relative spatial positions of amino acids in this system may not be a factor that will enhance our ability to predict the effects of mutations to nearby uncharacterized positions.

### Link between protein solubility and thermostability

Next, we explored the degree to which the 9 positions modified protein solubility. The loss of protein to inclusion body formation during expression and purification can be the result of altered folding thermodynamics, but it can also signal that the folding kinetics, or the native folding pathway, has been disrupted. Therefore, it is possible to produce a highly thermostable TEAS that nonetheless severely misfolds resulting in only modestly soluble protein ^25^.

We computed the most effective linear combination of mutations that together would drive changes in the in vivo solubility of TEAS heterologously expressed in *E. coli*. Solubilities were quantified as “high” if the concentration of eluted protein from Ni^+2^-NTA chromatography fell at or higher than the 75^th^ percentile (Fig. 5a). Given that all possible mutations were equally represented in the library, we computed the average mutation profile that produces high solubility rank (see Experimental Procedures), which is a simpler but conceptually similar dimensionality reduction analysis to the MNE approach ^26^. The result (Fig. 5b) represents the direction in M9 sequence space that predicts high protein solubility with the highest probability across the sequence population. Projection of the 512 mutant profiles onto this “preferred solubility profile” (blue distribution, Fig. 5c) compares well with the distribution arising from the 75^th^ percentile mutations (red distribution); that these two probability distributions have similar variances implicates only one significant mode of covariance, and, therefore, one mutational pattern that significantly influences protein solubility across the 9 positions.

**Figure 5.**
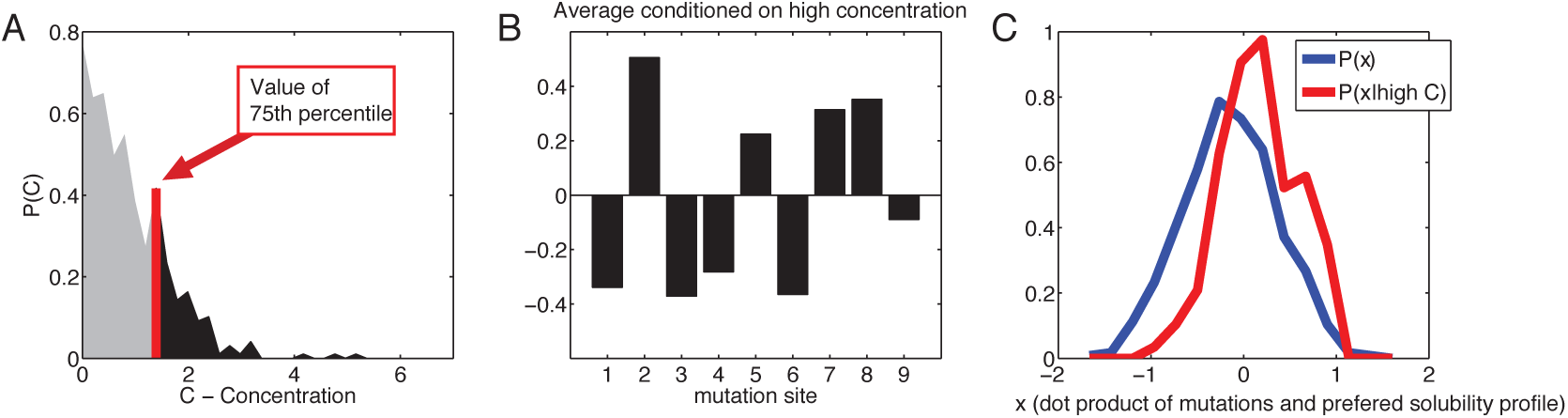
Average mutation conditioned on high concentration of soluble protein recovered from the M9 TEAS library. (A) Protein concentrations are plotted and the 75% percentile establishes the cutoff for “high solubility” mutants. (B) The preferred solubility profile is the linear combination of mutation frequencies at each position that most accurately predicts high solubility within the library. Y-axis represents each mutation’s contribution to high solubility. (C) The mutation profiles are projected onto the preferred solubility profile, where the blue curve is the probability of mutation for the entire dataset, and the red curve is the probability of mutation given high protein solubility. While their averages change, the variances of the two distributions do not significantly change, indicating there is likely only one direction of change across the protein concentration dataset and an analogous covariance analysis was not necessary.

The preferred solubility profile confirms that mutating positions that increase T_m_ (2, 5, 8) also improve solubility and that the T_m_ destabilizing positions (1, 3, 4, 6, 9) also reduce the probability of achieving high TEAS solubility (compare Figs. 4c and 5b). The exception to this correspondence is the position 7 mutation (I438T), which has almost no effect on T_m_s but provides robust improvements to TEAS solubilities, suggesting that sequence populations with Thr at position 7 (exchanging a methyl moiety for a hydroxyl group) may modify the folding kinetics and pathway while leaving the T_m_s of the folded proteins relatively unchanged.

### Non-additive pairwise mutational effects measured by MNE

The more nonlinear a system is, the more difficult it becomes to accurately predict outcomes of combined perturbations. In general, proteins are highly non-linear systems, and the structure and dynamic outputs of mutations tend to influence protein functions in complex ways. This phenomenon, often referred to as protein epistasis, has been assessed through modeling ^27^, measured directly using thermodynamic cycle analysis ^28,29^, and has been witnessed in real-time evolution of viral strains ^30^. Understanding protein epistatic behaviors is critical to predicting the functional consequences of sequence divergence within a protein family over time ^31^, and is essential for engineering proteins to behave like natural proteins ^32^. The importance of epistasis or non-additivity in our TPS system is suggested by the observation that the greatest changes in unfolding profiles is triggered by variation at position 4 with average shifts in T_m_s (compare Figs. 3c and 4b). This result implies that the effect of position 4 contextually depends on mutations at other locations. This indeed appears to be the case.

To characterize pairwise residue-residue coupling within the M9 library, we again applied MNE analyses, but now incorporating sequence populations encompassing pairs of mutations. Results for all pair-wise combinations of mutant populations are shown as black curves in Figure 6a (the first order MNE profiles are shown as well for comparison). Each of the mutant pairs exhibits statistically significant effects on thermal stabilities with similarly complex profiles as observed for the individual mutation analyses. Comparison of these profiles and the profiles computed assuming the effects of two mutations can be combined linearly quantifies the synergy, or the degree of non-additivity, within the mutant pair population. In Fig. 6a, we combined the MNE profiles from the two positions analyzed independently (top row) and overlaid them in red with their respective pair-wise MNE profiles. If there is no synergy, the two profiles will match, indicating that the pair-wise effect of two simultaneous mutations (25% of library mutants) in the background of all other changes at the other 7 positions (75% of library mutants) on the thermostability profiles are linear combinations of their individual effects. If, however, the two profiles do not match, then this indicates that there is synergy, quantified as the magnitude of the difference between these two vectors (Fig. 6b-yellow shaded area). Matrices of these quantities, shown by ||*v_ij_* − *v_i_* − *v_j_*||^2^ give the degree of non-additivity between pairs of mutations. Here, positions 2 and 8, in general, display the most synergy with other positions, indicating that their overall effects on the thermofluor curves are the most sensitive to sequence context (Fig. 6c).

**Figure 6.**
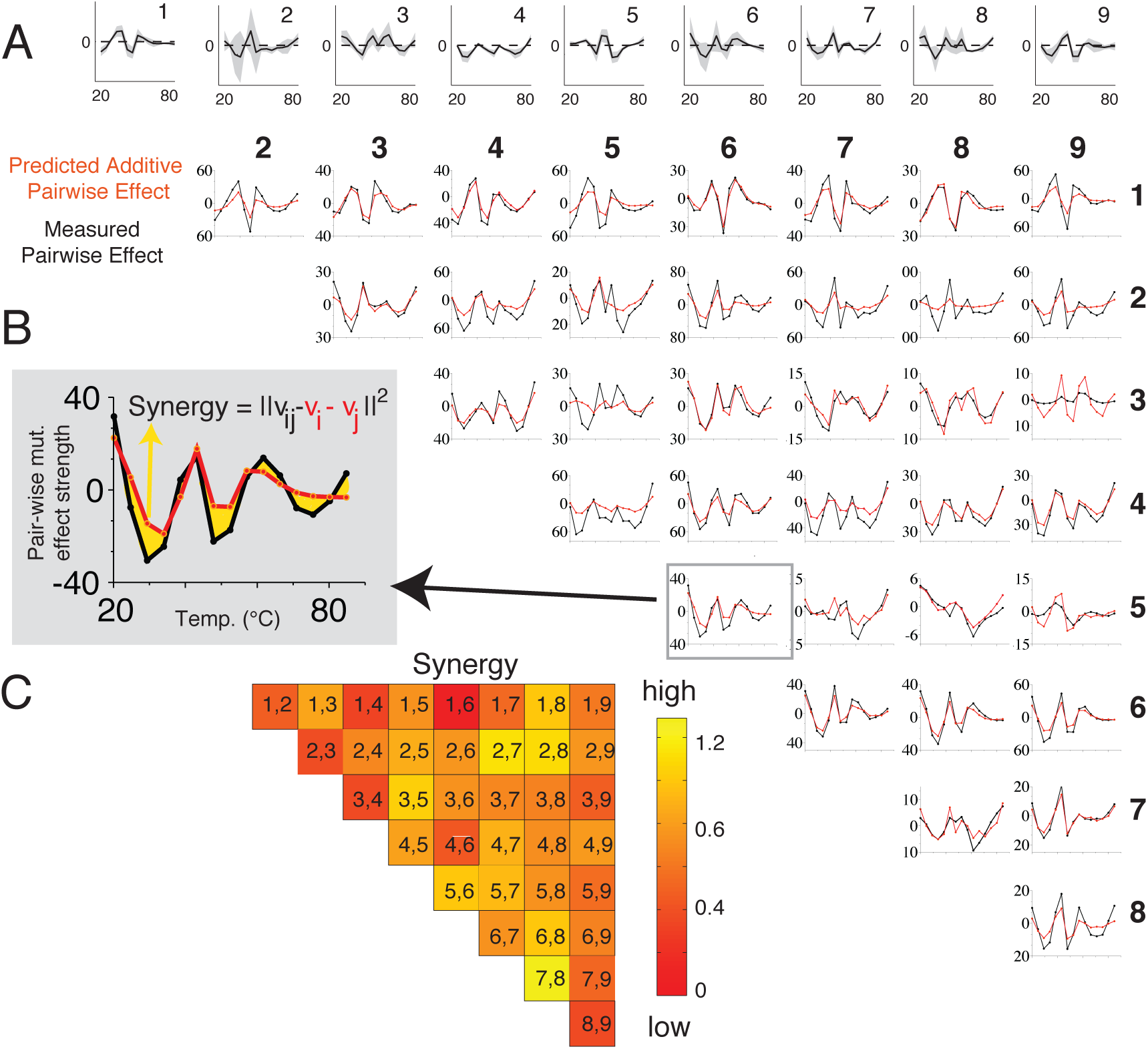
Second order MNE analysis reveals synergy between residues. (A) MNE profiles of second order effects from the 36 pairs of sites are shown in black and compared to the two first order MNE profiles that have been first scaled by the MNE norm. The difference between the two curves quantifies the temperature dependence of the double mutation nonlinearity, or the epistasis between the two positions. All MNE filters were found to be significant. (B) Synergy calculated from comparisons of expected MNE effects (red curve) and the calculated MNE effects (black curve). The magnitude of synergy (yellow shaded region) is exemplified using the pair 5:6. C. Matrix of mutational effect synergy between the 36 pairs.

Having quantified nonlinarity across the entire thermofluor curve, we then turned our attention to investigating the nonlinearity specific to the Tm, as this feature is a main target to improve enzyme robustness within increasing temperature extremes. We also quantified the effect of pairs of mutations on T_m_ by comparing these results to T_m_ shifts expected for independent contributions from each mutation in a given pair, averaged over the rest of the library. In the majority of cases, T_m_ shifts induced by pairs of mutations (red bars in Figure 7) were smaller than those expected from additive contributions from each mutation to T_m_ shifts. These suppressive effects are observed even when both positions shift the T_m_ in the same direction. For example, positions 1,3 and 6 are all destabilizing, but when paired together, their destabilizing effects are less than expected from a linear combination of their individual effects. The same is true for T_m_-stabilizing positions, i.e., 2, 5 and 8. Taken together, these pairwise analyses reveal that residue-residue mutations synergistically dampen changes to T_m_, especially when the additive effect is expected to be large (Fig. 7b).

**Figure 7.**
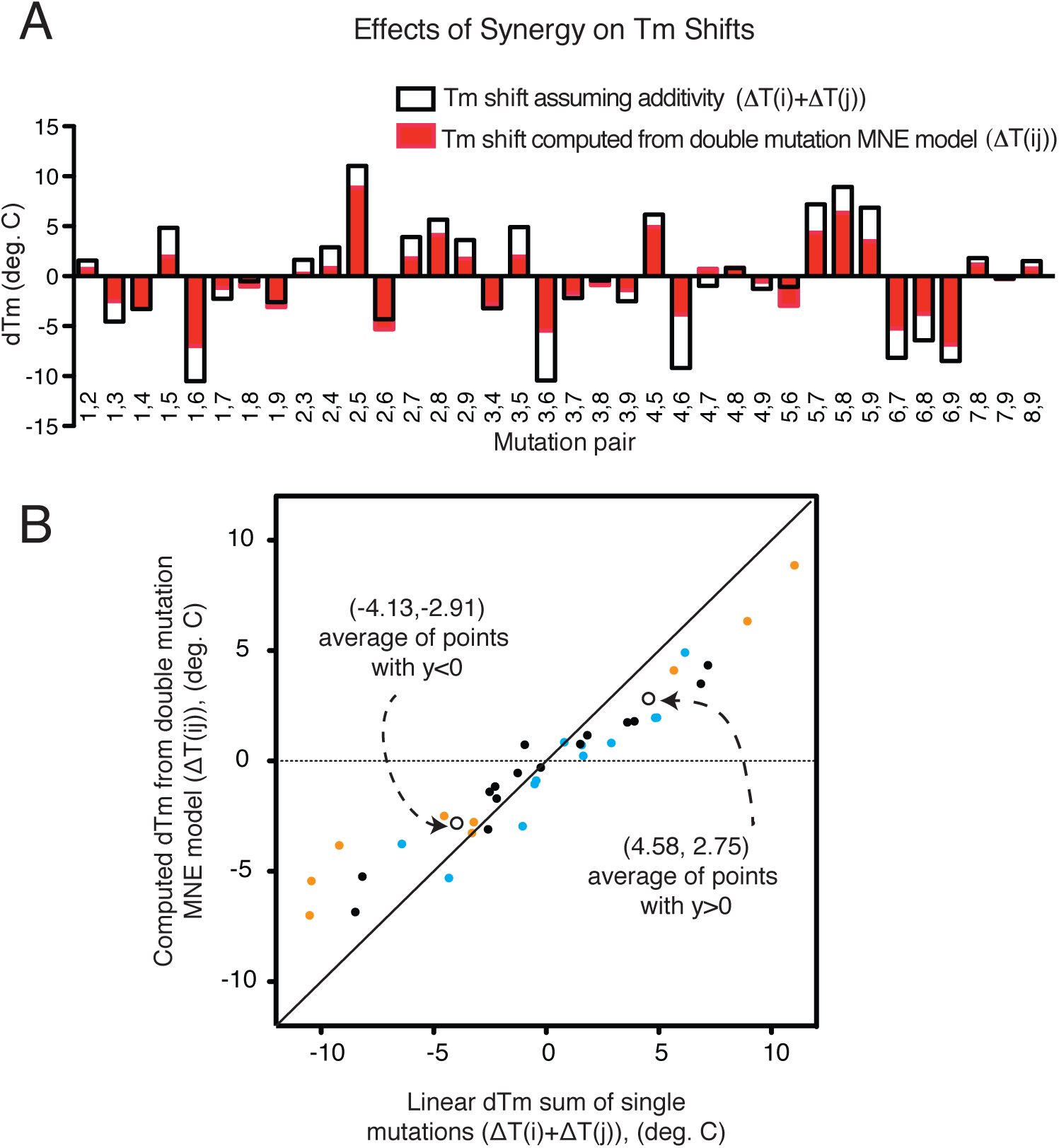
Effects of Synergy between pairs on T_m_ shifts. (A) Black outlined bars represent effects on the T_m_s that are predicted if the individual effects (from figure 4b) are additive. The red bars are the amounts of T_m_ shifts generated from the measured pairwise effects. The differences between the two bars demonstrate the role of synergy for each pair. (See text for discussion of calculations). (B) Scatter plot of pairwise data from dT_m_ calculated assuming additive pairwise effect (x axis) vs. dT_m_ from the measured pairwise effect (Y axis). Black diagonal line represents equality. On average, pairs of mutations cooperate to shift the T_m_ towards the WT value, even when both positions are stabilizing or destabilizing. Orange and blue points represent pairs that shift Tm in the same or opposite direction, respectively. Black points are pairs including positions 7 and 9, which minimally affect Tm. Open circles are averages which show the deviation from equality.

## Discussion

We set out to assess the rational evolvability of the TPS fold with the goal of generating enhanced robustness to thermal extremes. Using large biochemical datasets measuring thermal unfolding, enzyme solubility and catalytic product specificity, we found that several principles of this system were quite amenable to rational engineering. First, a cohort of thermostable mutants with Tm increases of up to ∼11degrees C is accessible within a relatively few number of mutations overall. We also identified three positions driving this elevated thermostability phenotype, which have effects so penetrant that they are stereotypical even in the background of mutations at the other positions. The utility of the MNE methodology is also highlighted by this work, as these population level measurements are made possible by these computational approaches tailor made for such large datasets.

Secondly, we found that T_m_ and catalytic specificity are uncorrelated, thus providing greater flexibility towards the independent optimization of these two crucial biophysical features of the enzyme. It will be interesting for future research to determine whether covariance exists between patterns of product profile variance and features of thermostablility profile variance beyond the T_m_. Third, we found a mutational profile that correlates with high solubility, thereby enabling accurate prediction of the relative solubility of any mutant in this library. Our reference system of the TEAS M9 library shows the TPS family will likely remain robust in the face of dynamic climate changes, as wide robustness-enhancing variances in product output, thermostability features and solubility are accessible within the same sequence space.

On the other hand, we also discovered that positions that stabilize or destabilize the fold are not spatially clustered in any strongly predictable way. Most importantly, as was previously demonstrated with TEAS catalytic profiles ^12^, the combined Tm effects of two mutations displays strong nonlinearity, which is on average dampening. These properties make it difficult to predictably hit particular sets of phenotypic targets through mutations. Taken together, these findings indicate that while protein engineering promises to reveal many mutational strategies for increasing specialized enzyme, and therefore crop, robustness, the inherent nonlinearity and non-predictability of these systems will continue to present challenges to the efficiency of these efforts.

Moving forward, we must be cautiously optimistic and emphasize the importance of diversified strategies both in scientific research and in the public policy arenas when dealing with the impacts of climate change. Indeed, failure to facilitate every available approach may have a significant adverse impacts on geopolitical stability. We hope that the coordinated efforts of researchers, including bioengineers/breeders can better inform policy makers and aid in the timely implementation of initiatives to address the demands of a growing population facing increasing environmental challenges.

## Online Methods

### Expression and purification of M9 library proteins

Expression and purification of library proteins was performed as previously described ^33^. Mutants were expressed in BL21(DE3) cells in 5 ml of Terrific broth (TB) growth media with kanamycin until cultures reached OD600≥0.8. Protein expression was induced by addition of IPTG to 0.1⃠mM followed by growth with shaking at 20⃠°C. for 5⃠h. Pellets from harvested cell cultures were re-suspended by adding 0.811ml of lysis buffer containing 1⃠mg⃠ml^-1^ lysozyme and 1⃠mM EDTA directly to frozen pellets followed by shaking at room temperature. Proteins were purified using 96-well, 800 uL, 25 to 30 um melt blown polypropylene filter plates (Whatman 7700-2804) and Ni-NTA superflow resin (Qiagen 30410). The N-terminal Histidine tag was left on the purified protein during the rest of the analysis and included a cleavage recognition site for either TEV or Thrombin protease. Protein purity and concentration were estimated by SDS-PAGE. Bands were quantitated using ImageJ ^34^ and compared to a 1 µM internal standard of WT TEAS protein.

For further details see Extended Experimental Procedures.

### Thermofluor assay and generation of unfolding phenotype neighbor joining tree

Thermal unfolding experiments were carried out in white 384-well plates in a LightCycler 480 (Roche Applied Science). Each well contained the protein in 25 mM Tris-HCl, pH 8.0, 250 mM NaCl, 50% glycerol (v/v), 2.5 mM β-mercaptoethanol and 125 mM imidazole and 10x SyproOrange Dye (Invitrogen). The plate was ramped from 20 to 85 °C with 10 data points acquired per °C. The dye fluorescence was detected at 580 nm using the dynamic integration mode (max integration time, 999 ms). Light Cycler→ 480 II software computed the global minimum of the first derivatives of the raw fluorescence versus temperature to obtain the T_m_ where possible. The temperature vs. fluorescence raw data were averaged (n=4) and used to generate a neighbor joining tree: temperature values were divided into 100 bins using a spline function and fluorescence values were normalized using L_2_ normalization where the sum of squared values equals 1. This approach preserves the shapes of the curves while placing each trace on the same relative fluorescence scale, (y^2^_1_+y^2^_2_+…+y100^2^)0.5. A 512X 512 Euclidean distance matrix was then calculated and used as input to generate a neighbor-joining tree in Mega 5.2 ^35^. Clades were designated heuristically by iteratively grouping the closest branches and measuring the average distances between these groups to look for smaller groups that could be combined without significantly changing the average intragroup distance.

### Maximum Noise Entropy analysis of Thermofluor data

## Overview and definitions

We used the Maximum Noise Entropy (MNE) ^18,19^ approach to find statistically significant associations of the enzyme fold stability and the mutational background of the protein. The end goal of this analysis is to compute a vector, also sometimes referred to as a “feature” or “filter”, that describes the direction along which two populations of data can be most effectively separated. Specifically, we search for a feature of the unfolding curves *v_i_* that best allows us to predict whether a given experimentally observed curve *Fq*(*T*) is is associated with the presence or absence of a mutation at site *i*.

As a technique for determining these features from a dataset, the MNE method is designed to minimize the bias imposed by making assumptions on the nature of correlations between the enzyme’s amino acid sequence {*m*_i_} and the protein’s unfolding curve *Fq*(*T*). It does so by maximizing the model’s uncertainty about all the correlations between mutations and unfolding curves (the noise entropy) outside of a set of constraints derived from the experimental data. We compute the MNE feature for each mutation site *i* = 1, …,9, and also for each of the 36 possible combinations of pairs of mutations at sites *i* and *j*. For analysis, we discretized the unfolding curves by binning them into *n = 15* equally sized temperature bins centered at *T_µ_* (*µ* = 1, …, *n*) that span the range of temperatures that was probed experimentally.

Importantly, the magnitude of the resulting vectors *v_i_* (or *v_ij_*, in the case of mutation pair analysis at sites *i* and *j*) quantifies the degree of separability (technically, the Kullback-Leibler distance) between the distributions of curves associated with mutated and WT enzymes projected onto the vector *v*. Hence, for each site (or site-pair) the magnitude of this vector, which we denote as *b,* quantifies the impact of a given mutation on the temporal unfolding profile.. For presentation purposes, all *v*’s are normalized to 1 (*v* · *v* = 1) whereas all the information about the magnitude resides in *b* values shown in Figure 3.

## Estimation of T_m_ shifts from the MNE profiles

We compared the degree to which the observed changes in thermal unfolding profiles caused by mutation at a particular site or site-pair can be understood exclusively as shifts in T_m_, without modifications of the unfolding profile. Denoting the unfolding profile of the WT TEAS protein as *F*_*wt*_(*T*) and its derivative as *F′*_*wt*_(*T*), the shift in melting temperature caused by a mutation is

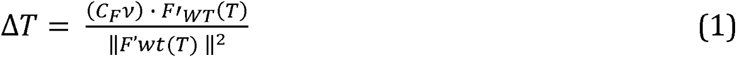

where *v* is the unnormalized MNE vector of the specific site or site-pair. The matrix *C_F_* is the covariance matrix of all unfolding curves in the dataset. Detailed derivation of Eq. (1) is provided in Extended Experimental Procedures

## Author Contributions

C.L. and P.E.O. expressed and purified the mutant protein library. C.M.C and I.R.F. generated protein thermostability and concentration datasets. J.A. and T.O.S. designed performed computational analyses. H.J.K. wrote scripts for computing biochemical phylogeny. C.M.C., J.A, T.O.S. and J.P.N wrote the paper.

## Acknowledgments

This work was supported by an Innovation Grant from The Salk Institute for Biological Studies and the National Science Foundation under award number EEC-0813570 to J.P.N. and IIS-1254123 to T.O.S. C.M.C. was funded, in part, by the National Institutes of Health T32 Cancer Training Grant, 5 T32 CA009370. We acknowledge support for P.E.O. from the Biotechnology and Biological Sciences Research Council (BBSRC) Institute Strategic Program Grants BB/J004561/1 (Understanding and Exploiting Plant and Microbial Secondary Metabolism). We declare no competing financial interests related to this work. J.P.N. is an investigator with the Howard Hughes Medical Institute.

